# Resolving the Taxonomic Status of the Marbled Toad (Bufonidae: *Incilius marmoreus*): 2RAD-based Phylogeography Including an Isolated Population in Veracruz, Mexico

**DOI:** 10.1101/2024.11.23.624927

**Authors:** Kenneth Wang, Todd W. Pierson, Joseph R. Mendelson

**Author notes:** Send reprint requests to JRM.

## Abstract

*Incilius marmoreus* inhabits an extensive range along the Pacific Coast of Mexico and a smaller allopatric region in the State of Veracruz, exhibiting an unusual distribution among herpetofauna. In 1901, Günther classified the Pacific coastal toads *Bufo argillaceus* and *B. lateralis* as conspecific with *Incilius [Bufo] marmoreus,* which has its type locality in Veracruz. Here, we adopt a multidisciplinary approach to reevaluate the phylogeography and taxonomy of *I. marmoreus* by gathering and analyzing morphological data and conducting phylogenetic and population genetic analyses from genome-wide SNP data. Our results uphold the current taxonomy by concurring with Günther. Our phylogenetic and population genetic analyses suggest that *I. marmoreus* from Veracruz are closely related to those from Oaxaca whilst coalescent analyses recovered a north–south split along the Pacific Coast estimated to have occurred ∼0.86 Mya followed by a shallow east–west split in the southern lineage that separates the Pacific coastal populations and the allopatric population in Veracruz ∼0.33 Mya. This species displays marked morphological and genetic diversity throughout its range, but this variation appears to be consistent with gene flow across contiguous populations rather than the existence of independent evolutionary lineages. The processes leading to the geographic isolation of the population on the coast of Veracruz remain uncertain, but we hypothesize that climatic and vegetation changes in the Late Pleistocene may have played a role.

**JOURNAL PUBLICATION STATEMENT:** This article has been published in *Ichthyology & Herpetology*: https://doi.org/10.1643/h2024107.

## INTRODUCTION

Species are fundamental units of biology and systematics (Mayr, 1982; de Queiroz, 2007). Thus, systematic studies often concern species delimitation—the determination of whether different populations represent one or more distinct species (Liu et al., 2021; Pereyra et al., 2021; Rivera et al., 2022; Dufresnes, 2025). Despite general agreement that species are independently evolving metapopulation lineages (de Queiroz, 2005, 2011), investigators differ in their interpretation of which species criteria (e.g., reciprocal monophyly, ecological differentiation, phenetic diagnosability, reproductive incompatibility) are most salient to diagnose species. Consequently, research in the field can sometimes yield variable, unstable, or uncertain results due to theoretical disagreements over species concepts and philosophy (Frost and Kluge, 1994; de Queiroz, 2007; Hillis, 2019), methodological concerns such as the appropriateness of genetic loci used in phylogenetic analysis (Chan et al., 2022), geographic inclusiveness of sampling with respect to hybridization zones and areas of sympatry (Myers et al., 2020; Marshall et al., 2021), and discordance among and within morphological and molecular datasets (e.g., Toews and Brelsford, 2012; Denton et al., 2014). Discordant patterns may be the result of phenomena such as incomplete lineage sorting (Wang et al., 2018), poor resolution from sampled loci (Dell’Ampio et al., 2014), or reticulate evolutionary patterns produced by hybridization and introgression (Ruane et al., 2014; Johnson et al., 2015; Myers et al., 2020; Pyron et al., 2020; Burbrink et al., 2021; Chambers et al., 2023).

Due to its complex geography and tremendous biodiversity, Mexico provides many opportunities to study the biogeography and systematics of herpetofauna—of which there are >1,000 species of non-avian reptiles (52% endemic to the country) and >400 species of amphibians (70% endemic to the country; Suazo-Ortuño et al., 2023). Phylogeographic studies of these taxa shed light not only upon their focal species, but also upon the shared evolutionary histories of other organisms in the region (e.g., Bryson et al., 2011; Ríos-Muñoz and Navarro□Sigüenza, 2012; Suárez-Atilano et al., 2014). Broadly speaking, the Mexican Transition Zone is the area where the Nearctic and Neotropical realms grade into each other (Halffter, 1987; Morrone, 2020), and it includes five distinct mountainous features: the Trans-Mexican Volcanic Belt, Sierra Madre Occidental, Sierra Madre Oriental, Sierra Madre del Sur, and the Chiapan–Guatemalan Highlands. The former three major mountain ranges and their associated highlands form the interior Central Mexican Highlands, which is sandwiched by coastal lowlands; the southerly Chiapan–Guatemalan Highlands, are separated from the other ranges by the lowland Isthmus of Tehuantepec (see below). The formation of these mountainous features was complex, with orogeny mostly occurring or finishing during the Miocene (Mastretta-Yanes et al., 2015). For this reason, many phylogeographic studies attribute recent vicariant speciation (e.g., in the Pleistocene) to climatic changes and older events to geological activity (Bryson et al., 2012; Mastretta-Yanes et al., 2015; Schramm et al., 2021). Nonetheless, the four magmatic episodes that gave rise to the relatively young Trans-Mexican Volcanic Belt were not complete until ∼3 Mya in the Plio-Pleistocene (Ferrari et al., 2012).

In southern Mexico, the Isthmus of Tehuantepec separates the Sierra Madre del Sur from the Chiapan–Guatemalan Highlands, connecting the Veracruzan and Pacific Lowlands biogeographic provinces across a contiguous, lowland ∼200 km east–west zone (Morrone et al., 2017). Some geological and biological evidence supports the existence of a Pliocene–Pleistocene trans-isthmian seaway (Durham et al., 1955; Sullivan et al., 2000; Mulcahy et al., 2006; Daza et al., 2010; Hardy et al., 2013), which may have further isolated highland species, although this remains contentious (Durham et al., 1955; Barrier et al., 1998; Campbell, 1999; Mulcahy et al., 2006). Regardless, the Isthmus of Tehuantepec is a well-recognized vicariant barrier between a diverse variety of highland taxa including mammals, birds, and herpetofauna (Duellman, 1960; Sullivan et al., 1997; Arellano et al., 2005; Weir et al., 2008; Castoe et al., 2009; Zamudio-Beltrán et al., 2020). Although not as well-documented, it is also a biogeographic break between lowland taxa found in the tropical moist forest on the northern (Gulf of Mexico) side and tropical dry forest and thornscrub on the southern (Pacific) side. For example, sister species of *Coleonyx* geckos (Eublepharidae) are found on either side, with the abrupt transition from the northern wet tropical forest habitat of the Gulf to the southern dry tropical forests of the Pacific acting as an ecological barrier to gene flow (Butler et al., 2023).

Here, we focus on uncovering the phylogeography of a lowland species with allopatric populations found on the Pacific and Gulf coasts (but not continuously across the intervening lowlands in the Isthmus of Tehuantepec)—the Marbled Toad (*Incilius marmoreus*). Originally described as *Bufo marmoreus* Wiegmann, 1833, we apply the genus name *Incilius* Cope, 1863 following recommendations suggested by morphology and molecular data (Frost et al., 2006; Frost et al., 2009; Mendelson et al., 2011). *Incilius* spp. are commonly referred to as “Mesoamerican Toads”, although some species extend beyond the traditional boundaries of Mesoamerica—south into the Ecuador (e.g., *I. coniferus*) and north into the southern United States (e.g., *I. nebulifer* and *I. alvarius*). *Incilius marmoreus* is endemic to Mexico and is distributed predominately in xeric to subhumid and deciduous habitat along the Pacific Coast, from the southern tip of Sonora to near Nachig, Chiapas. Interestingly, the type specimens of *I. marmoreus* originate from the city and state of Veracruz, which borders the Gulf of Mexico, and are thus rather disjunct from the remainder of the distribution (Fig. 1A). These Veracruzan records appear to be primarily associated with palm savannah and scrub habitat (Rzedowski, 1994). Records of *I. marmoreus* from northern Hidalgo (Lemos-Espinal and Dixon, 2016) were determined to instead represent *I. nebulifer* (JRM unpubl. data), and a single record in Durango (Lemos-Espinal et al., 2019) remains unconfirmed and is separated by approximately 220 kilometers from the populations on the Pacific. From our examination of museum specimens (see Methods), this species does not exist in the Balsas Depression, but specimens of the morphologically similar *I. perplexus* (Taylor, 1943) that occurs there are often confused with *I. marmoreus* (see following paragraph). Thus, verified records of *I. marmoreus* are known from two allopatric regions: a broad expanse of the Pacific Coast and a small area in Veracruz. While broad distributions of animal species along the subhumid Pacific Coast of Mexico are common, such a distribution with an allopatric population in coastal Veracruz is not (an example is noted in the Discussion). *I. marmoreus* appears to be abundant throughout much of its range and is currently considered a species of Least Concern (IUCN SSC Amphibian Specialist Group, 2020), but modeling suggests that it might face a loss of distribution in the next ∼60 years from climate change (Garcia et al., 2014).

**Figure 1.**
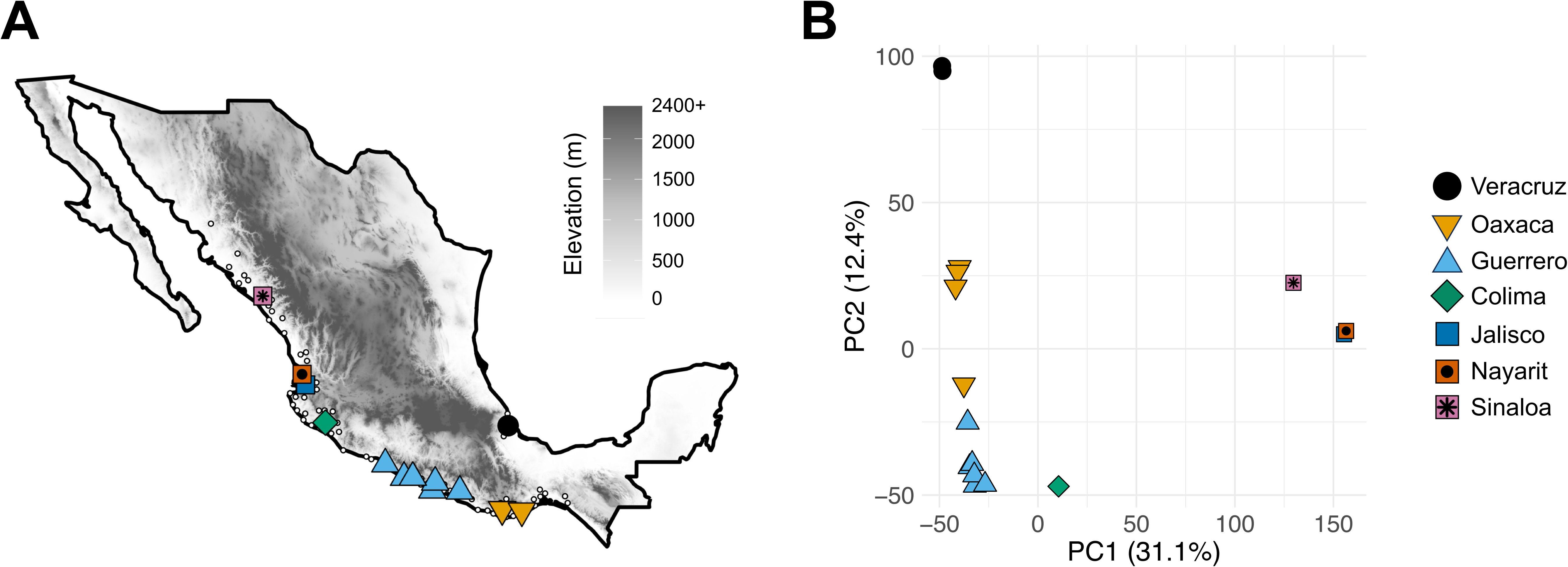
(A) A map of Mexico showing the distribution of the Marbled Toad (*Incilius marmoreus*). Small white dots represent research-grade and unobscured iNaturalist hosted occurrence points downloaded from GBIF; we used the R package spThin version v0.2.0 (Aiello-Lammens et al., 2015) to spatially thin records at a threshold of 25 km. Larger solid symbols represent locations from which we generated 2RAD sequencing data. Identifications were verified through dorsal patterning and crest morphology. Shading indicates elevation. (B) The first two principal components (PC) axes from a principal components analysis (PCA) of 2RAD data.

*Incilius marmoreus* is sexually dimorphic; as males mature, they lose their juvenile marbled pattern and develop a solid dorsal coloration that ranges from greenish yellow to yellow-gold, whilst females develop a brown coloration and retain their dorsal pattern (van der Heiden, 2022). Along the Pacific Coast, the morphologically similar *I. perplexus* is also sexually dichromatic, but it can be distinguished from *I. marmoreus*. In females, the dorsal pattern of *I. perplexus* is sprawled and rounded whereas *I. perplexus* bears more triangular blotches. In males, it is more difficult to differentiate between species (e.g., from specimens that have lost their patterning). However, although the parotoid glands of *I. marmoreus* can be either triangular or rounded, the parotoid glands of *I. perplexus* are always rounded and generally “pill-shaped”. Along the Pacific Coast, *I. marmoreus* may also distinguished from sympatric *I. mazatlanensis* (Taylor, 1940 “1939”) by the absence or strong reduction of parietal crests, which are present and well-developed in *I. mazatlanensis*. In Veracruz, *I. marmoreus* may be distinguished from sympatric *I. valliceps* (Wiegmann, 1833) in the same manner.

According to Frost (2024), three taxa have been previously synonymized with *Incilius marmoreus* Wiegmann, 1833*. Bufo argillaceus* Cope, 1868 and *Bufo lateralis* Werner, 1894 were described along the Pacific Coast, with the type localities in Colima and Oaxaca, respectively. However, type specimens corresponding to these taxa were deemed to be conspecific with *B. [Incilius] marmoreus* by Günther (1901). Until now, this recommendation to consider the allopatric (and morphologically similar) *I. marmoreus* in the Pacific and Gulf lowlands as conspecific has not received further attention. In addition, *Bufo eiteli* Ahl, 1927 (with the type locality in Veracruz) was later synonymized with *B. [Incilius] marmoreus* by Gorham (1974). Because previous work on *Incilius* spp. has shown that many allopatric montane populations represent distinct species (reviewed by Mendelson et al., 2011), reexamination of the two allopatric populations of *I. marmoreus* is warranted. Here, we adopt a multidisciplinary approach involving both morphological comparison of museum specimens and phylogenetic and population genetic analyses of genome-wide single nucleotide polymorphisms (SNPs) to test divergence under the general lineage species concept (de Queiroz, 1998, 2007) and evaluate alternative phylogeographic hypotheses. We also comment on differences in interspecific relationships from our results compared to those from prior phylogenetic studies (Mendelson et al., 2011).

## MATERIALS AND METHODS

### Morphological data collection and analysis

We obtained EtOH-preserved specimens of *I. marmoreus* from the following museums: AMNH, CAS, KU, LACM, MCZ, MVZ, TCWC, UCM, UMMZ, UTA, and UTEP (museum codes per Sabaj et al., 2023). These included specimens collected from Veracruz (and deposited in the AMNH and UTA collections) and from across the remainder of the geographic range of the species. After verifying identifications, we measured a total of n = 171 specimens of *I. marmoreus*. Of the 126 males, 29 were from Veracruz; of the 45 females, 7 were from Veracruz. We confirmed sex through examination of nuptial excrescences on the first finger (present in males, absent in females) and corroborated these determinations using differences in dorsal color patterns (van der Heiden, 2022).

Following Duellman (1970) and Mendelson (1994), we collected the following measurements, rounded to the nearest 0.1mm, via a digital calipers: SVL (snout–vent length; the distance from the rostral tip to the posterior end of the ischium), TIB (tibial length), FT (foot length), HL (head length), HW (head width), EYE (diameter of the bony orbit), EN (eye–nostril length), TYMP (tympanum diameter, across the major axis if elliptical), PARL (longest parotoid measurement parallel to the sagittal plane), and PARW (longest parotoid measurement parallel to the coronal plane). We analyzed these data in three ways. First, we used Welch’s t-tests (α = 0.05) to evaluate whether the average SVL differed between animals from Veracruz and those from the Pacific Coast, separating males and females into two separate analyses. Second, we log-transformed the measurements and used a series of linear models to test for regional differences in the averages of other traits and for differences in how these traits scale with SVL; to do this, we again separated males and females into different analyses and fit models using the function “ln()” in R v4.3.1 (R Core Team, 2023). We evaluated statistical significance (α = 0.05; Bonferonni adjusted α = 0.0056) of interaction terms using Type-III sums of squares with function “Anova()” in the package “car” v3.1-2 (Fox and Weisberg, 2019). If the interaction term was not significant, we refitted the models without the interaction and evaluated statistical significance of the main effect (i.e., region) using Type-II sums of squares as described above. Third, we conducted principal component analyses (PCA) through singular value decomposition (Wall et al., 2003) using the “prcomp()” function in R v4.3.1 (R Core Team, 2023). We separated males and females into different analyses, included all log-transformed morphological measurements, and plotted the first two axes using the package “ggplot2” v3.5.1 (Wickham, 2016), with separate ellipses denoting 95% confidence intervals surrounding points from Veracruz and those from all other locations.

### 2RAD library preparation and sequencing

To generate genome-wide SNP data, we used a reduced-representation genome sequencing method. These methods, which include Restriction site Associated DNA sequencing (RADseq) and its many variants, sequence a subset of orthologous loci across genomes of many individuals and have become popular tools to investigate phylogenetic and population genetic questions in non-model organisms like amphibians and reptiles (e.g., Pante et al., 2015; McCartney-Melstad et al., 2018; Liu et al., 2020; Scott et al., 2020). Here, we used 2RAD (Bayona-Vásquez et al., 2019), which is similar to dual-digest RADseq (ddRAD) (Peterson et al., 2012) but offers several advantages such as better utility with low-concentration DNA samples and lower startup costs.

We prepared 2RAD libraries for 16 *I, marmoreus* (including 2 from Veracruz), 5 *I. perplexus*, 1 *I. canaliferus*, 5 *I. mazatlanensis*, 1 *I. valliceps*, and 1 *Rhinella horribilis*. We extracted DNA using a Qiagen DNeasy Blood and Tissue kit and quantified extracts using a Qubit 4.0 Fluorometer. We then prepared 2RAD libraries (Bayona-Vásquez et al., 2019) with 50 ng of input DNA and using the enzymes XbaI and EcoRI-HF. We conducted individual PCRs, pooled PCR products by volume, and size-selected this pool for 450–550 bp on a Pippin Prep. We then pooled these libraries with others from unrelated projected for paired-end 150-bp (PE150) sequencing on an Illumina NovaSeq 6000.

### 2RAD data assembly and analysis

We assembled demultiplexed data in ipyrad v0.9.94 (Eaton and Overcast, 2020), with the following parameters: reads trimmed to 130 bp, a clustering threshold of 0.85, a minimum read depth of 6 reads per locus, and a minimum of 4 samples per locus. We then used this assembly in the following phylogenomic and population genomic analyses.

First, we estimated a maximum-likelihood phylogeny from an alignment of all 2RAD loci using the program RAxML v8.2.12 (Stamatakis, 2014) and through the analysis toolkit in ipyrad v0.9.94 (Eaton and Overcast, 2020). We used a GTRGAMMA nucleotide substitution model and inferred 100 rapid bootstrap replicates followed by a full maximum-likelihood search. We rooted this phylogeny with the outgroup *R. horribilis*, and we plotted all bootstrap replicates in a DensiTree-like (Bouckaert, 2010) visualization using the package “phangorn” v2.11.1 (Schliep, 2011) in R v4.3.1 (R Core Team, 2023). Second, we used the species-tree inference program tetrad v.0.9.14, which is based upon the SVDQuartets algorithm (Chifman and Kubatko, 2014), to infer a species tree from SNPs. We used our full assembly, all quartets, and 100 bootstrap replicates, and we again rooted with the outgroup and plotted all replicates. Third, we estimated a distance-based phylogenetic network using the Neighbor-Net algorithm (Bryant and Moulton 2004) as implemented in the package “phangorn” (Schliep, 2011; Schliep et al., 2017) in R v4.3.1 (R Core Team, 2023) and using a pairwise Hamming distance matrix constructed from one random SNP per locus.

Fourth, we conducted a principal components analysis (PCA) of *I. marmoreus* samples using the analysis toolkit in ipyrad v0.9.94 (Eaton and Overcast, 2020). We first filtered our assembly to: 1) include only ingroup samples (i.e., *I. marmoreus*); 2) include only SNPs present in at least 80% of samples; 3) include only SNPs with a minor allele frequency (MAF) ≥ 0.2; and 4) impute missing data with *k-*means clustering and *k* = 6. We plotted the first two PC axes in R v4.3.1 (R Core Team, 2023). Fifth, we used the Bayesian clustering program Structure v2.3.4 (Pritchard et al., 2000) to evaluate population genetic structure, and we likewise conducted this analysis through the analysis toolkit in ipyrad v0.9.94 (Eaton and Overcast, 2020). We first filtered our assembly to: 1) include only ingroup samples (i.e., *I. marmoreus*); 2) include only SNPs present in at least 80% of samples; 3) include only SNPs with a MAF ≥ 0.05; and 4) include only one SNP per locus. We then conducted 10 replicate runs of 300,000 Markov chain Monte Carlo steps (with 30,000 burn-in steps) for *k* = 1–7. We removed replicates that did not converge (i.e., by setting max_var_multiple = 100) and visually examined a plot of ΔK (Evanno et al., 2005) to assess the “optimal” number of clusters. With the aid of CLUMPP (Jakobsson and Rosenberg, 2007), we then plotted the average of replicates for two strongly supported values of *k* overlayed on a map.

Finally, we used the program Bayesian Phylogenetics & Phylogeography (BPP; Rannala and Yang, 2003; Yang, 2015) through the analysis toolkit in ipyrad v0.9.94 (Eaton and Overcast, 2020) to estimate divergence times and effective population sizes under a multi-species coalescent model (i.e., the A00 analysis), modeling our analyses after Butler et al. (2023). We used only samples of *I. marmoreus*, and we conducted these analyses for two alternative scenarios: 1) one in which we modeled coalescence between Veracruz samples and all others; and 2) one in which we modeled coalescence between Veracruz and the remainder of southern samples (see Results), then coalescence between these groups and the northern samples. For each scenario, we assigned an inverse-gamma prior (3, 0.002) for the population size parameter θ and an inverse-gamma prior (2, 0.0015) for τ_0_, the divergence time of the root. We conducted two independent runs for each scenario, with each run using 100 random loci and 120,000 generations (with 20,000 discarded as burn-in). We then transformed our parameter estimates into units of time and effective population size using a gamma distribution of mutation rates bounded by values estimated for frogs (7.76 x 10^-10^ – 1.53 x 10^-9^ mutations per site per year (Crawford, 2003; Sun et al., 2015) and generation times of 0.8, 2, or 3 years (i.e., the minimum ages at reproduction reported for *Incilius*; Oliveira et al., 2017). We visually assessed Markov chain Monte Carlo (MCMC) mixing and convergence in Tracer v1.7.2 (Rambaut et al., 2018).

## RESULTS

### Morphological analyses

Male *I. marmoreus* from Veracruz (mean ± SD = 48.9 ± 2.9 mm) were smaller than those from the Pacific Coast (57.0 ± 4.2 mm) (*t* = -11.75, df = 66.82*, P* < 0.001). Likewise, females from Veracruz (52.6 ± 5.2 mm) were smaller than those from the Pacific Coast (61.5 ± 4.9 mm) (*t* = - 4.20, df = 8.06, *P* = 0.003). In males, we found a significant interaction between SVL and region (i.e., Veracruz vs. Pacific Coast) for HW (Table 1). For male toads from Veracruz, we found that HW was narrower at small SVLs but increased at a greater rate with SVL. We also found a significant interaction between SVL and region for EYE; smaller toads from Veracruz had larger eyes, and eye size increased at a more gradual rate with SVL. However, neither of these differences were significant after a Bonferroni correction for multiple comparisons (Table 1). We also found significant differences in the intercept (i.e., mean values) for EN and TYMP; both these differences were significant even after correction for multiple comparisons (Table 1). Males from Veracruz had proportionately larger EN and smaller TYMP. In females, we found a significant interaction between SVL and region (i.e., Veracruz vs. Pacific Coast) for PARL (Table 2). For female toads from Veracruz, we found that PARL was shorter at small SVL but increased at a greater rate with SVL. However, this difference was not significant after correction for multiple comparisons. We also found a significant difference in intercept (i.e., mean values) for TIB (Table 2); females from Veracruz had smaller TIB than those from elsewhere. This difference was not significant after correction for multiple comparisons (Table 2).

**Table 1.** Results from a series of linear models for morphological measurements of male Marbled Toads (*Incilius marmoreus*). We modeled each morphological variable as a function of snout–vent length (SVL) and its interaction with region (i.e., Veracruz vs. Pacific Coast). We used an analysis-of-variance (ANOVA) with Type III Sums of Squares to evaluate statistical significance. If the interaction term was significant, we do not report the main effect. If the interaction term was not significant, we refit the model without the interaction and used a Type-II sums of squares to evaluate the main effect of region. Here, we indicate statistical significance (α = 0.05) with an asterisk, and statistical significance after a Bonferroni correction (adjusted α = 0.0056) with two asterisks. All values rounded to three decimal places. TIB = tibial length; FT = foot length; HL = head length; HW = head width; EYE = eye diameter; EN = eye-nostril length; TYMP = tympanum diameter; PARL = parotoid length; PARW = parotoid width.

|  | SS (Type III) | SS (Type II) | F-value | Pr(>F) |
| --- | --- | --- | --- | --- |
| TIB slopes | 0.002 |  | 1.285 | 0.259 |
| TIB intercepts |  | 0.001 | 0.523 | 0.471 |
| FT slopes | 0.004 |  | 0.403 | 0.527 |
| FT intercepts |  | 0.000 | 0.000 | 0.988 |
| HL slopes | 0.003 |  | 1.806 | 0.182 |
| HL intercepts |  | 0.000 | 0.007 | 0.935 |
| HW slopes | 0.013 |  | 4.590 | 0.034* |
| HW intercepts | — | — | — | — |
| EYE slopes | 0.014 |  | 4.108 | 0.045* |
| EYE intercepts | — | — | — | — |
| EN slopes | 0.005 |  | 1.936 | 0.167 |
| EN intercepts |  | 0.043 | 18.242 | 0.000** |
| TYMP slopes | 0.023 |  | 2.124 | 0.148 |
| TYMP intercepts |  | 0.095 | 8.559 | 0.004** |
| PARL slopes | 0.049 |  | 1.742 | 0.189 |
| PARL intercepts |  | 0.066 | 2.326 | 0.130 |
| PARW slopes | 0.000 |  | 0.000 | 0.991 |
| PARW intercepts |  | 0.005 | 0.208 | 0.650 |

**Table 2.** Results from a series of linear models for morphological measurements of female Marbled Toads (*Incilius marmoreus*). We modeled each morphological variable as a function of snout–vent length (SVL) and its interaction with region (i.e., Veracruz vs. Pacific Coast). We used an analysis-of-variance (ANOVA) with Type III Sums of Squares to evaluate statistical significance. If the interaction term was significant, we do not report the main effect. If the interaction term was not significant, we refit the model without the interaction and used a Type II Sums of Squares to evaluate the main effect of region. Here, we indicate statistical significance (α = 0.05) with an asterisk, and statistical significance after a Bonferroni correction (adjusted α = 0.0056) with two asterisks. All values rounded to three decimal places. Abbreviations are provided in the text and Table 1 caption.

|  | SS (Type III) | SS (Type II) | F-value | Pr(>F) |
| --- | --- | --- | --- | --- |
| TIB slopes | 0.002 |  | 1.082 | 0.304 |
| TIB intercepts |  | 0.007 | 4.097 | 0.049* |
| FT slopes | 0.018 |  | 3.896 | 0.056 |
| FT intercepts |  | 0.010 | 1.880 | 0.178 |
| HL slopes | 0.008 |  | 2.703 | 0.108 |
| HL intercepts |  | 0.000 | 0.017 | 0.897 |
| HW slopes | 0.010 |  | 4.070 | 0.050 |
| HW intercepts |  | 0.000 | 0.020 | 0.889 |
| EYE slopes | 0.000 |  | 0.001 | 0.981 |
| EYE intercepts |  | 0.015 | 3.880 | 0.055 |
| EN slopes | 0.003 |  | 0.956 | 0.334 |
| EN intercepts |  | 0.005 | 2.021 | 0.163 |
| TYMP slopes | 0.014 |  | 1.074 | 0.306 |
| TYMP intercepts |  | 0.030 | 2.252 | 0.141 |
| PARL slopes | 0.113 |  | 5.439 | 0.025* |
| PARL intercepts | — | — | — | — |
| PARW slopes | 0.007 |  | 0.315 | 0.578 |
| PARW intercepts |  | 0.003 | 0.138 | 0.712 |

In both PCAs of morphological traits, we found that the first principal component (PC) axis mostly separated animals from Veracruz from those from all other locations, but 95% confidence ellipses overlapped considerably (Fig. 2). For both males and females, SVL had the highest loading for this axis, with animals from Veracruz being smaller than those from elsewhere (Tables 3, 4). Other measurements with high loadings on this first axis were TIB, HL, HW, and EN. In the case of males, the second PC axis emphasized variation in size and shape of the parotoid glands, with strong positive loadings on PARL and PARW. The third PC axis assigned strong positive loadings on EYE and PARW. In females, the second PC axis mostly explained eye-related measurements, with strong positive loadings on EYE and EN. The third PC axis emphasized parotoid variation, with strong positive loadings on PARL and PARW.

**Figure 2.**
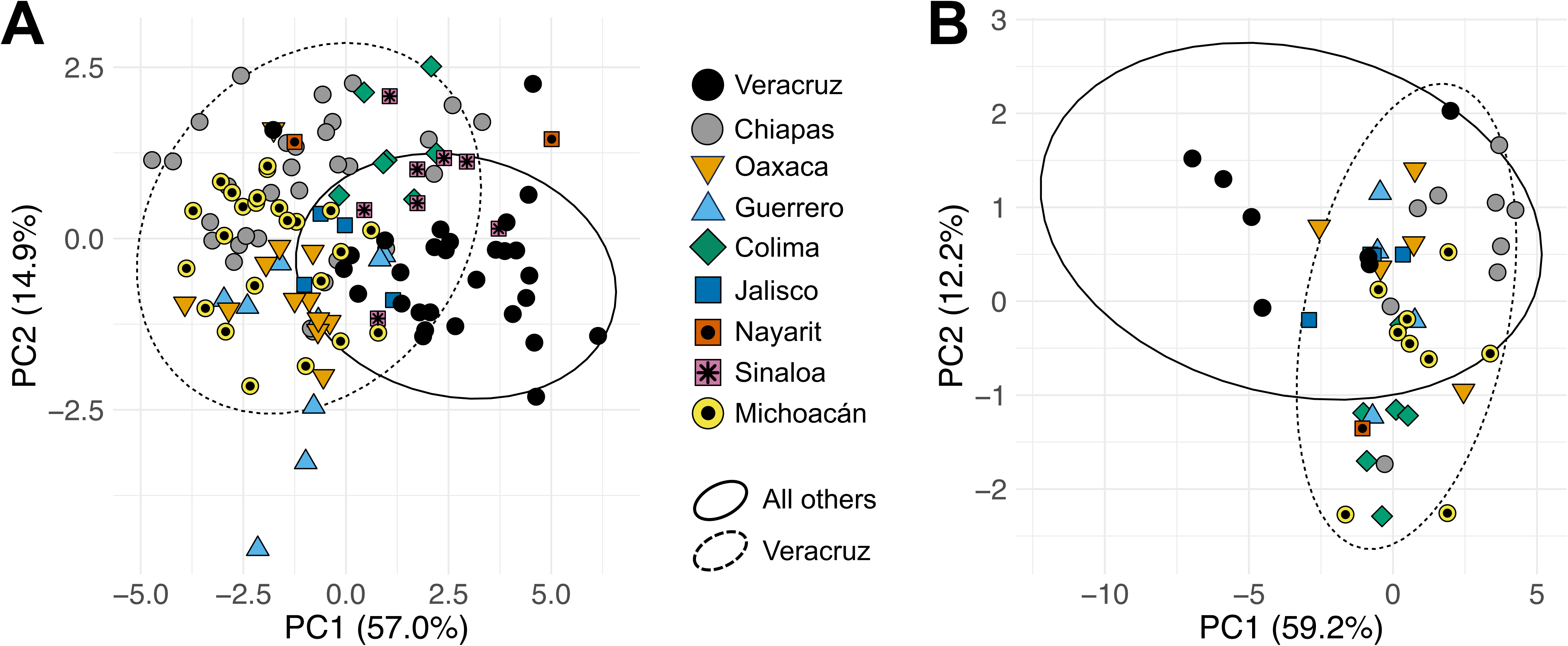
(A) The first two principal components (PC) axes from a principal components analysis (PCA) of morphological data collected from male Marbled Toads (*Incilius marmoreus*) from across their geographic distribution. (B) The first two PC axes from an identical analysis of female *I. marmoreus*. In both, point colors and shapes reflect the collection location, and ellipses show 95% confidence intervals around two groups: toads from Veracruz and toads from the Pacific Coast.

**Table 3.** Results from a principal components analysis (PCA) of morphological measurements from male Marbled Toads (*Incilius marmoreus*). Here, we report results from all principal components (PC) axes up to 80% cumulative variance explained. Abbreviations are provided in the text and Table 1 caption.

|  | PC1 | PC2 | PC3 |
| --- | --- | --- | --- |
| Proportion of Variance | 0.5697 | 0.1493 | 0.0874 |
| Cumulative Variance | 0.5697 | 0.7190 | 0.8064 |
| Eigenvalue | 5.6974 | 1.4928 | 0.8741 |
| SVL | -0.3973 | -0.0113 | -0.0953 |
| TIB | -0.3724 | 0.0647 | -0.2269 |
| FT | -0.3231 | 0.0287 | -0.1367 |
| HL | -0.3782 | -0.0956 | -0.0276 |
| HW | -0.3605 | 0.0136 | -0.1008 |
| EYE | -0.2475 | -0.4432 | 0.5608 |
| EN | -0.3354 | -0.3412 | 0.3037 |
| TYMP | -0.2999 | 0.1012 | -0.4738 |
| PARL | -0.2100 | 0.5434 | 0.2451 |
| PARW | -0.1354 | 0.6059 | 0.4674 |

**Table 4.** Results from a principal components analysis (PCA) of morphological measurements from female Marbled Toads (*Incilius marmoreus*). Here, we report results from all principal components (PC) axes up to 80% cumulative variance explained. Abbreviations are provided in the text and Table 1 caption.

|  | PC1 | PC2 | PC3 |
| --- | --- | --- | --- |
| Proportion of Variance | 0.5917 | 0.1219 | 0.0865 |
| Cumulative Variance | 0.5917 | 0.7135 | 0.8001 |
| Eigenvalue | 5.9168 | 1.2186 | 0.8652 |
| SVL | 0.3714 | -0.0923 | -0.1983 |
| TIB | 0.3613 | -0.2457 | -0.1049 |
| FT | 0.3266 | -0.1941 | 0.0317 |
| HL | 0.3586 | -0.0241 | -0.2519 |
| HW | 0.3647 | -0.0075 | -0.2648 |
| EYE | 0.2125 | 0.6961 | -0.0612 |
| EN | 0.3243 | 0.4577 | -0.1033 |
| TYMP | 0.2490 | -0.4209 | 0.0359 |
| PARL | 0.3042 | -0.0782 | 0.4629 |
| PARW | 0.2416 | 0.1245 | 0.7650 |

### Phylogenomic and population genomic analyses

We obtained 1.8–39.0 million PE150 reads per sample (median = 14.6 million), and our final assembly included > 306,000 SNPs across > 75,000 loci. Our maximum-likelihood phylogenetic analysis recovered *I. marmoreus* as sister to *I. perplexus,* and these two taxa together were then sister to *I. canaliferus* (RAxML bootstrap support [b.s.] = 100). Together, these three taxa were sister to *I. mazatlanensis* + *I. valliceps* (b.s. = 100; Fig. 3A). We recovered all these interspecific relationships with high bootstrap support, and the topologies of the species-tree inference was concordant (Fig. 3B). The distance-based phylogenetic network showed superficially conflicting results (e.g., *I. valliceps* clustering with *I. canaliferus* and the outgroup; Fig. 3C). However, because the network is unrooted, we encourage caution in the interpretation of the species-level topology it suggests. Within *I. marmoreus*, each of these analyses suggested a northern Pacific-Coast clade consisting of samples from Sinaloa, Nayarit, and Jalisco (b.s. = 100) and a southern Pacific-Coast clade consisting of samples from Guerrero, Oaxaca, and Veracruz (b.s. = 100); we recovered the sample from Colima as sister to the southern clade in the maximum-likelihood and species-tree analyses, and we recovered it as intermediate between the northern and southern clades in the phylogenetic network (Fig. 3). In all three analyses, samples from Veracruz were nested within samples from Oaxaca (Fig. 3).

**Figure 3.**
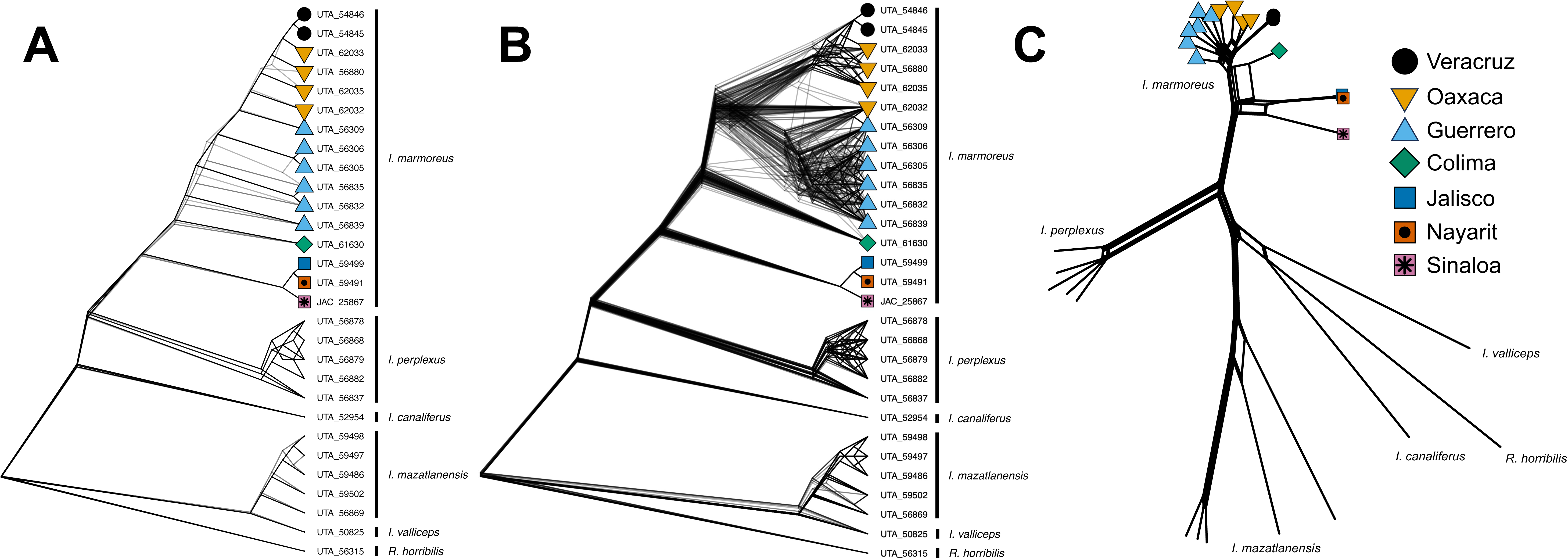
(A) DensiTree-like plots of RAxML rapid bootstrap replicates from a maximum-likelihood analysis of loci from 2RAD data, showing relationships within the Marbled Toad (*Incilius marmoreus*) and between relatives; (B) a similar plot showing bootstrap replicates from a tetrad species-tree analysis of SNPs from the same dataset; (C) a Neighbor-Net phylogenetic network inferred from one random SNP per locus from the same dataset. In all, point colors and shapes reflect the collection location within Mexico.

The results of the PCA on the molecular data were consistent with our phylogenetic analyses. The first PC axis explained 31.1% of variation and separated the northernmost samples (i.e., from Sinaloa, Nayarit, and Jalisco) from the more southern samples, with the single sample from Colima being somewhat intermediate, but closer to the southern samples (Fig. 1B). The second PC axis explained 12.4% of variation and differentiated samples within the southern clade, with those from Veracruz and Guerrero widely separated and those from Oaxaca at intermediate positions (Fig. 1B). The Structure results provided corroborating evidence for these same broad patterns. We found highest support for *k* = 2 and *k* = 4. At *k* = 2, the two clusters corresponded with the northern and southern Pacific-Coast clades described above, and the sample from Colima showed intermediate assignment values (Fig. 4A). At *k* = 4, the samples from Sinaloa were placed in a third cluster (i.e., distinct from others in the northern clade), and the fourth cluster had highest assignment probabilities in the samples from Veracruz, with those from Guerrero and Oaxaca showing increasing assignment probabilities to this cluster moving from west to east (Fig. 4B).

**Figure 4.**
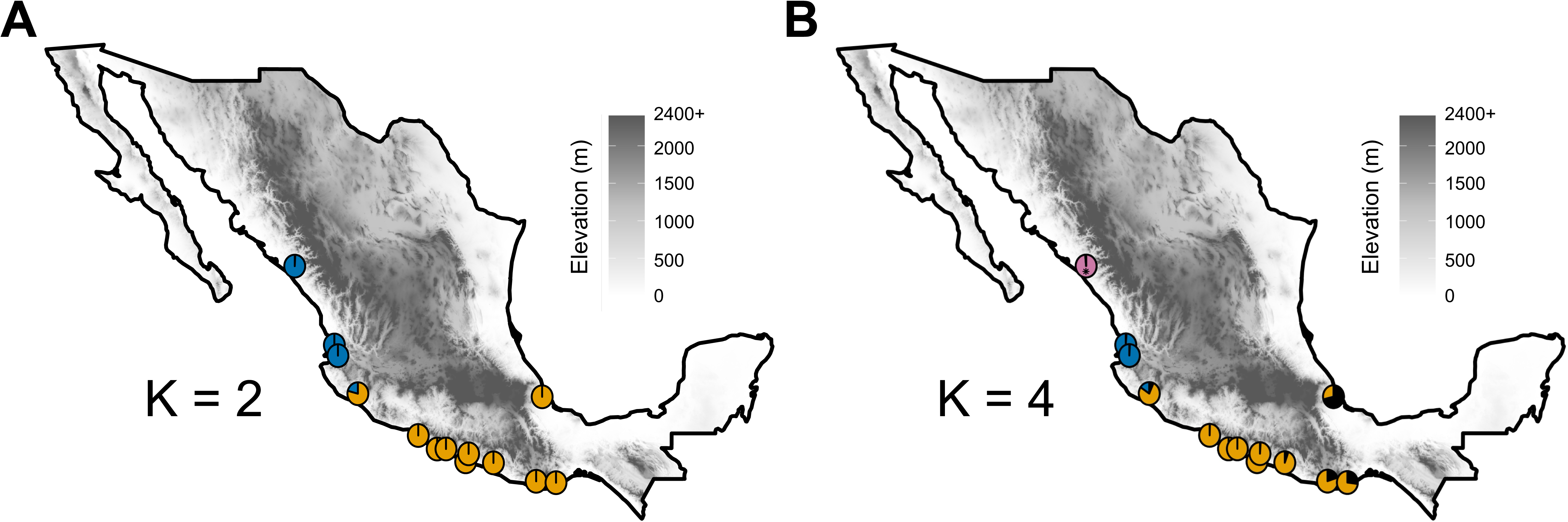
Genetic clustering results for the Marbled Toad (*Incilius marmoreus*), inferred using SNPs from 2RAD data and with the program Structure. Plots show results from (A) *k* = 2 and (B) *k* = 4. In both plots, some pie charts represent averages of assignments from more than one individual sample from the same collection site. Shading indicates elevation.

In our coalescent analyses in BPP, we found relatively concordant results across our two modeled scenarios. In the first scenario, we estimated a median coalescent time of 0.33 Mya (95% credible interval [CI] = 0.19–0.56) for Veracruz (median N_e_ with generation time of 0.8 years [N_e,0.8_] = 0.11 M [95% CI = 0.06–0.19]; N_e,2.0_ = 0.04 M [0.02–0.08]; N_e,3.0_ = 0.03 M [0.02–0.05]) and all other populations (N_e,0.8_ = 2.03 M [1.15–3.47]; N_e,2.0_ = 0.81 M [0.46–1.39]; N_e,3.0_ = 0.54 M [0.31–0.92]) (Fig. 5A). In the second scenario, we estimated a coalescent time of 0.31 Mya [0.17–0.53] for Veracruz (N_e,0.8_ = 0.10 M [0.05–0.17]; N_e,2.0_ = 0.04 M [0.02–0.07]; N_e,3.0_ = 0.03 M [0.01–0.05]) and the southern Pacific-Coast populations (N_e,0.8_ = 0.82 M [0.44–1.46]; N_e,2.0_ = 0.33 M [0.18–0.58]; N_e,3.0_ = 0.22 M [0.12–0.39]) and a coalescent time of 0.86 Mya [0.51–1.41] between these groups (N_e,0.8_ = 0.47 M [0.21–0.95]; N_e,2.0_ = 0.19 M [0.08–0.38]; N_e,3.0_ = 0.13 M [0.06–0.25]) and northern Pacific-Coast samples (N_e,0.8_ = 0.71 M [0.42–1.18]; N_e,2.0_ = 0.28 M [0.17–0.47]; N_e,3.0_ = 0.19 M [0.11–0.31]) (Fig. 5B).

**Figure 5.**
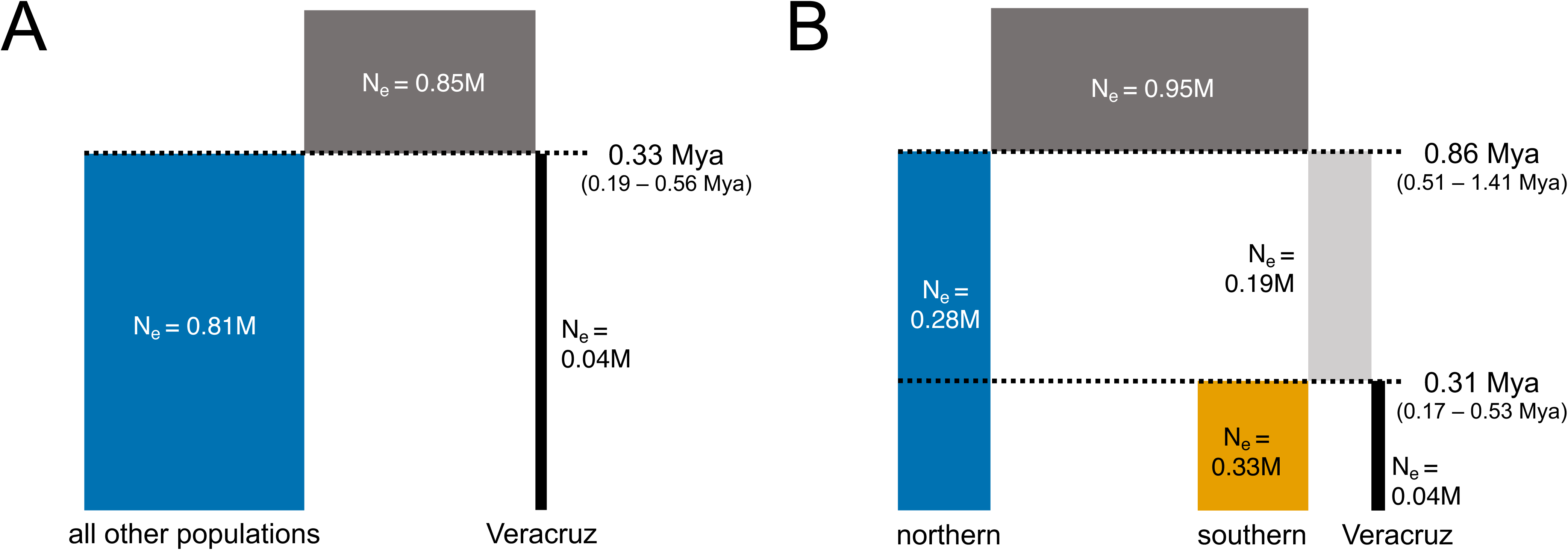
Results from BPP coalescent analyses with two alternative demographic scenarios: (A) one in which we estimated a date for the coalescence between Veracruz samples and all others; and (B) one in which we estimated a date for the coalescence of Veracruz and the remainder of southern samples (see Fig. 4) and a second date for the coalescence between these two and the northern samples. Within each figure, bar widths are proportional to estimated effective population sizes (N_e_), and bar heights are proportional to estimated coalescence dates. For simplicity, the effective population sizes show only estimates from transformation using a generation size of two years.

## DISCUSSION

Our investigation revealed that *I. marmoreus* displays substantial morphological and genetic variability throughout its range. Based on our sampling, populations in Veracruz have smaller SVL and some allometric differences from those along the Pacific Coast, but analyses of genome-wide SNP data suggest that they are phylogenetically nested within populations from Oaxaca—the most geographically proximate sampling sites from across the Isthmus of Tehuantepec. Our inclusion of both genomic and morphological data facilitated a robust analysis of this group, and we argue that our results are consistent with recognition of a single species. Below, we further discuss our results and their implications for the phylogeography and systematics of *Incilius*.

From gross visual inspection, we did not observe any geographic cline in the morphological proportions or color pattern along the Pacific Coast, nor could we identify any diagnostic trait for distinguishing Pacific toads from those in Veracruz. However, our morphometric analyses suggest that *I. marmoreus* from Veracruz have a smaller average SVL. Additionally, we found some evidence that male *I. marmoreus* from Veracruz exhibit head widths that scale more steeply in relation to SVL, eye diameters that scale more gradually with SVL, proportionately longer eye–nostril measurements, and proportionately smaller tympana compared to their Pacific counterparts. We also found some evidence that female *I. marmoreus* from Veracruz exhibit parotoid lengths that scale more steeply in relation to SVL and proportionately shorter tibiae.

We acknowledge that these morphological analyses have several limitations. First, they include only adult specimens. Additionally, statistical analyses implicitly assume linear relationships between SVL and other morphological measurements, but some traits may scale otherwise. Furthermore, statistical power for the female comparisons (e.g., only seven females from Veracruz) may have been impaired since our sampling was rather male-biased, as is often the case for collections of bufonid toads (Mendelson 1994). Perhaps most importantly, we note that all but two (UTA A-54845, A-54846) measured toads from Veracruz were drawn from a single museum series collected from the type locality of the city of Veracruz, reducing our confidence in the generalizability of the differences we measured.

Both our maximum-likelihood phylogeny and species tree differed in species-level topology from the summary hypothesis tree presented in Mendelson et al. (2011). We found strong support for a sister relationship between *I. marmoreus* and *I. perplexus*, with *I. canaliferus* being in-turn sister to *I. marmoreus* + *I. perplexus*; in contrast, Mendelson et al. (2011) hypothesized a sister relationship between *I. canaliferus* and *I. perplexus*, with *I. marmoreus* being sister to *I. canaliferus* + *I. perplexus*. Our topology is more concordant with morphological evidence, for *I. perplexus* and *I. marmoreus* resemble each other in gross morphology and pattern whilst *I. canaliferus* is readily distinguishable from them (and in fact, all other *Incilius* spp.) by its distinctively long parotoid glands.

Within *I. marmoreus,* our genome-wide SNP data were consistent with isolation-by-distance along the Pacific Coast, Phylogenetic analyses demonstrate a northernmost and southernmost clade with high support whilst PCA and Structure analyses corroborates this primary axis of phylogeographic variation. However, it is worth noting that these phylogenetic analyses (and our coalescent analyses in BPP) do not explicitly incorporate gene flow—a limitation that could result in a biased inference of topology and demographic parameters. Perhaps unexpectedly, we discovered that samples from Veracruz were nested within samples from Oaxaca in phylogenetic analyses (but clustered separately in the PCA); thus, we found evidence that toads from the southern end of their Pacific distribution (Guerrero and Oaxaca) are more closely related to toads from Veracruz than they are to toads from the contiguous distribution farther north along the Pacific Coast (Jalisco, Nayarit, Colima). The estimated coalescent timing ∼0.31 or 0.33 Mya between toads in Veracruz and those nearest them on the Pacific Coast is considerably later than the hypothesized ∼2 Mya Pliocene–Pleistocene marine introgression into the Isthmus of Tehuantepec (Maldonado-Koerdell, 1964; Sullivan et al., 2000) and also prevents us from newly proposing a hypothesis of anthropogenic transportation (Ardelean et al., 2020; Becerra-Valdivia and Higham, 2020).

Nonetheless, exactly what series of events may explain the current allopatric distribution of *I. marmoreus* remains uncertain. With an extensive distribution along the western coast of Mexico and a smaller allopatric distribution in Veracruz, Mexico, the geographic distribution of *Incilius marmoreus* is rather unusual, but not unprecedented among herpetofauna. For example, the crotaline viper *Agkistrodon bilineatus* covers a somewhat similar distribution, with those in Veracruz having been described as *A. bilineatus lemosespinali* from a single specimen (Smith and Chiszar, 2001). However, the provenance of that specimen and validity of that taxon have been questioned (Campbell and Lamar, 2004; Bryson and Mendoza-Quijano, 2007; Porras et al., 2013). Drawing from Gloyd and Conant (1990), Bryson and Mendoza-Quijano (2007) hypothesized that a narrow band of seasonally dry forest—running from northern Guerrero to Veracruz—might have acted as a dispersal corridor for *Agkistrodon*. We find this to be an unlikely explanation for the biogeography of *I. marmoreus* for two reasons. First, unlike other groups with similar distributions (e.g., *Ctenosaura* iguanas; Zarza et al., 2019), these toads appear not to currently live in the interior lowlands (i.e., the Balsas Depression) nearest this hypothetical corridor. Second, our data showed that *I. marmoreus* in Veracruz were most closely related to those in southern Oaxaca—not those from Guerrero. This seemingly incongruous pattern also was found by Zarza et al. (2019) where they reported that the southernmost populations of the Pacific coastal iguanid lizard *Ctenosaura pectinata* were most closely related to its sister species, *C. acanthura* from the Veracruzan region, than to the more northerly conspecific populations along the Pacific Coast. Unlike our results, there is ample evidence to recognize the Veracruzan *C. acanthura* as a species distinct from the Pacific coastal *C. pectinata*.

Instead, we suggest an alternative hypothesis. We propose that *I. marmoreus* diversified throughout the Pacific Coast of Mexico and then dispersed eastward into Veracruz via the Isthmus of Tehuantepec. Afterwards, relatively recent isthmian climatic and vegetative changes, resulting in a wet Gulf forest and dry Pacific forest, may have ecologically isolated the Veracruz population with the development of unsuitable wet-forest habitat on the northern side of the isthmus. Today, the eastern terminus of the Trans-Mexican Volcanic Belt may prevent *I. marmoreus* from expanding its distribution further north along the Gulf coast, limiting its northern extent to near the town of Palma Sola, Veracruz. Indeed, this eastern terminus is a biogeographic barrier between congeners *I. valliceps* and *I. nebulifer* (Mulcahy et al., 2006) and has been implicated in the diversification of many other taxa, such as several fish species (Mateos, 2005; Huidobro et al., 2006).

We recognize *I. marmoreus* as a single species exhibiting geographic variation—a conclusion drawn from both morphological and genomic data. Consequently, we concur with Günther (1901) that the names *Bufo argillaceus* and *Bufo lateralis* represent junior synonyms of *I. marmoreus*. We were able to further corroborate this synonym by examining photographs of the syntypes or holotypes of *B. marmoreus, B. argillaceus, and B. lateralis.* Furthermore, we were able to verify the later synonymy of *B. eiteli* with *B. marmoreus* by Gorham (1974) in the same manner. Although the populations in Veracruz remain disjunct and are apparently somewhat dwarfed, they share a very recent evolutionary history with populations on the Pacific Coast and are not diagnosable by any discrete morphological diagnostic characters. Thus, under the general lineage concept of species, we do not recognize the *I. marmoreus* in Veracruz as representing an independently evolving metapopulational lineage, and hence not as a distinct species (de Queiroz, 1998; de Queiroz, 2007). Our study is not the first to recognize populations of distinctly diminutive animals as conspecific with “normal-sized” populations. For example, Hoza et al. (2023) used genomic data to argue against the recognition of dwarf *Phrynosoma hernandesi* that is geographically isolated in the San Luis Valley, Colorado, USA, as a distinct species. Likewise, along the Pacific Coast, gene flow, lack of morphological distinguishability, and lack of evidence of ecological divergence do not support the evolutionary distinctiveness of the northern and southern metapopulations.

Finally, we note that the basic natural history of *I. marmoreus* remains nebulous. For example, *I. marmoreus* has been reported to deposit eggs singly (Blair, 1972). However, this behavior is highly unusual amongst bufonids and otherwise unknown within *Incilius* (but see image by Campuzano León, 2016). The veracity of this reported behavior deserves corroboration, as single egg-laying behavior has been wrongly attributed to other bufonids, such as *Anaxyrus debilis* (Girard, 1854) and *Anaxyrus retiformis* (Sanders and Smith, 1951). In the case of these two species, eggs are, in truth, laid in short and fragile strings that break easily, consequently appearing as if they have been laid singly (Ferguson and Lowe, 1969; Altig and McDiarmid, 2015). We encourage future researchers to describe and document additional aspects of this species’ life history, behavior, and ecology.

## Supporting information

Figure 1 Alt-Text

Figure 2 Alt-Text

Figure 3 Alt-Text

Figure 4 Alt-Text

Figure 5 Alt-Text

Table 1 Summary

Table 2 Summary

Table 3 Summary

Table 4 Summary

## DATA ACCESSIBILITY

All raw sequence reads are available through the NCBI SRA (PRJNA1173256). The assembled 2RAD dataset, morphological data, and code for all analyses are available through Zenodo (10.5281/zenodo.17393182).

## ACKNOWLEDGEMENTS

We thank Greg Pauly, Ana Motta, Carol Spencer, David Kizirian, Emily Braker, Greg Schneider, Lee Fitzgerald, Neftali Camacho, Stevie Kennedy-Gold, and Vicky Zhuang for facilitating specimen and tissue loans. Daniel G. Mulcahy, Frank Tillack, Gassner Georg, Kevin de Queiroz, Mark-Oliver Rödel, and Schweiger Silke for providing photos of the syntypes and holotypes associated with Incilius marmoreus and its junior synonyms I. argillaceus, I. lateralis, I. Eiteli. Marco Arturo Puente-Zozaya for the Spanish translation of the abstract, and Jacobo Reyes-Velasco for his significant comments and reviewal during the drafting process.

## APPENDIX 1 [Specimens Examined]

### Incilius marmoreus

Mexico: Chiapas: 2.7 mi [4.3 km] W Chiapa de Corzo (AMNH 173339–40), 18 mi [29.0 km] N Arriaga (AMNH 173341–42); 29.5 mi [47.5 km] NE Tapanatepec, Oaxaca (AMNH 173343–51); 21.6 mi [34.8 km] N Arriaga (AMNH 173352–54); 13.3 mi. [21.4 km] S. Tuxtla Gutierrez, along Chis. 28 (UCM 48536–41); 4.8 mi [7.7 km] S. Jct. Mexico 190 and road to La Concordia, along road to La Concordia (UCM 48544–50); 10.6 mi [17.1 km] S. Jct. Mexico 190 and road to La Concordia, along road to La Concordia (UCM 48551–56); 15.1 mi [24.3 km] S. Jct. Mexico 190 and road to La Concordia, along road to La Concordia (UCM 48557); 0.4 mi [0.6 km] NNE (by road on Mexico Highway 35) of Arriaga (UTEP 4714); 4.3 mi [7.0 km] N of Arriaga (UTEP 5843); 1.0 mi [1.6 km] N of Arriaga (UTEP 5927).

Mexico: Colima: 7.5 mi [12.1 km] SW Colima (LACM 36957, 59–70).

Mexico: Guerrero: 3.5 mi [5.6 km] S San Andres de la Cruz (KU 86676, 86679, 86681–82, 86684, 86688–89); Hwy 200 btwn Pie de la Ruesta turnoff and 15 mi [24 km] W (LACM 128416); .6 mi [1 km] NW Puerto Marquez above Hwy to airport (LACM 129899); 24 mi [39 km] (by Mex Hwy 200) WNW Coyuca de Benitez, 1.6 mi [2.6 km] by rd SE Alcholoa (LACM 129910–11); Acapulco (MCZ 1436); Chilpancingo (MCZ 17756).

Mexico: Jalisco: 5 mi [8 km] E Melaque on Hwy 80 (KU 95440–42, 45–46); Village of Casimiro Castillo on Hwy 30 (LACM 87827); 6 mi [10 km] NE of Chamela, La Huerta (UTEP 5860–61).

Mexico: Michoacan: 23.1 mi [37.2 km] WNW Caleta de Campos (TCWC 55735–39); 1 mi [1.6 km] NE of La Placita 50 ft [15 m] (UMMZ 104543, (1) specimen); NE of La Placita 75 ft [23 m] (UMMZ 104544, (1) specimen); 1 mi [1.6 km] NE of La Placita 75 ft [23 m] (UMMZ 104547); 0.5 mi [0.8 km] SW of La Placita (UMMZ 104548, (2) specimens); Ostula 500 ft [152 m] (UMMZ 104550, (5) specimens); near Pomaro along tributary to Rio Pasnori (UMMZ 104808, (15) specimens); near Rancho El Diezmo 4.5 mi [7.2 km] N of Coire 2920 ft [890 m] (UMMZ 104810, (2) specimens); Playa Azul (UMMZ 112790, (2) specimens).

Mexico: Nayarit: 21-25 mi [34-40 km] S. Acaponeta, 100’ +/- Hwy 15 (LACM 25403);

Compostela: 3 mi [5 km] SE of Sayulita (UTEP 5855–56).

Mexico: Oaxaca: Pochutla 150 (KU 58322, 24–25, 28–29, 31); Hwy 185 between Tehuantepec & Salina Cruz (LACM 8172–81, 65054).

Mexico: Sinaloa: 1 mi [1.6 km] ENE San Lorenzo (KU 47926, 30, 32–33, 39–42).

Mexico: Veracruz: Vera Cruz, western part of city (AMNH 12282–83, 86–89, 91–92, 94–96, 12302–03, 10–11, 13–14, 16, 20, 23, 26–27, 29, 36, 38); [city of] Vera Cruz (AMNH 13390–91, 99, 13404, 14, 25, 28, 32); town of Pochote, 2.3 rd mi [3.7 km] NW jct Hwy 140 at Tierra Colorada (UTA A-54845–46).

## LITERATURE CITED

Ahl, E. 1927 “1926”. Ueber neue oder seltene Froschlurche aus dem Zoologischen Museum Berlin. Sitzungsberichte der Gesellschaft Naturforschender Freunde zu Berlin 1926:111–117.

Aiello-Lammens, M. E., R. A. Boria, A. Radosavljevic, and B. Vilela. 2015. spThin: an R package for spatial thinning of species occurrence records for use in ecological niche models. Ecography 38:541–545.

Altig, R. I., and R. W. McDiarmid. 2015. Handbook of Larval Amphibians of the United States and Canada. Comstock Publishing Associates, Ithaca, New York.

Ardelean, C. F., L. Becerra-Valdivia, M. W. Pedersen, F. J. Herrera, A. S. Moreno-Mayar, E. M. R. Thomas, J. D. Ingversen, M. T. P. Gilbert, D. A. Hengst, J. C. Arroyo-Cabrales, E. J. Kennett, R. González-Reynoso, I. R. Becerra-González, J. A. Chatters … E. Willerslev. 2020. Evidence of human occupation in Mexico around the Last Glacial Maximum. Nature 584:87–92.

Arellano, E., F. X. González-Cozátl, and D. S. Rogers. 2005. Molecular systematics of Middle American Harvest Mice *Reithrodontomys* (Muridae), estimated from mitochondrial cytochrome b gene Sequences. Molecular Phylogenetics and Evolution 37:529–540.

Barrier, É., L. Velasquillo, M. Chávez, and R. Gaulon. 1998. Neotectonic evolution of the Isthmus of Tehuantepec (southeastern Mexico). Tectonophysics 287:77–96.

Bayona-Vásquez N. J., T. C. Glenn, T. J. Kieran, T. W. Pierson, S. L. Hoffberg, P. A. Scott, K. E. Bentley, J. W. Finger, S. Louha, N. Troendle, P. Diaz-Jaimes, R. Mauricio, and B. C. Faircloth. 2019. Adapterama III: Quadruple-indexed, double/triple-enzyme RADseq libraries (2RAD/3RAD). PeerJ 7:e7724.

Becerra-Valdivia, L., and T. Higham. 2020. The timing and effect of the earliest human arrivals in North America. Nature 584:93–97.

Blair, W.F. 1972. *Bufo* of North and Central America, p. 93–101. In: Evolution in the Genus Bufo. W. F. Blair (ed.). University of Texas Press, Austin, Texas.

Bouckaert, R. R. 2010. DensiTree: making sense of sets of phylogenetic trees. Bioinformatics 26:1372–1373.

Bryant, D., and V. Moulton. 2004. Neighbor-net: an agglomerative method for the construction of phylogenetic networks. Molecular Biology and Evolution 21:255–265.

Bryson, Jr. R. W., and F. Mendoza-Quijano. 2007. Cantils of Hidalgo and Veracruz, Mexico, with comments on the validity of *Agkistrodon bilineatus lemosespinali*. Journal of Herpetology 41:536–539.

Bryson, Jr., R. W., R.W. Murphy, A. Lathrop, and D. Lazcano-Villareal. 2011. Evolutionary drivers of phylogeographical diversity in the highlands of Mexico: a case study of the *Crotalus triseriatus* species group of montane rattlesnakes. Journal of Biogeography 38:697–710.

Bryson, Jr., R. W., U.O. García-Vázquez, and B.R. Riddle. 2012. Diversification in the Mexican horned lizard *Phrynosoma orbiculare* across a dynamic landscape. Molecular Phylogenetics and Evolution 62:87–96.

Burbrink, F. T., G. M. Gehara, A. D. McKelvy, and E. A. Myers. 2021. Resolving spatial complexities of hybridization in the context of the gray zone of speciation in North American ratsnakes (*Pantherophis obsoletus* complex), Evolution 75:260–277

Butler, B. O., L. L. Smith, and O. Flores-Villela. 2023 Phylogeography and taxonomy of *Coleonyx elegans* Gray 1845 (Squamata: Eublepharidae) in Mesoamerica: the Isthmus of Tehuantepec as an environmental barrier. Molecular Phylogenetics and Evolution 178:107632

Campbell, J. A. 1999. Distribution patterns of amphibians in Middle America, p. 111–209. In: Distribution Patterns of Amphibians: A Global Perspective. W.W. Duellman (ed.). John Hopkins University Press, Baltimore, Maryland.

Campbell, J. A., and W. L. Lamar. 2004. Venomous reptiles of the Western Hemisphere. Cornell University Press, Ithaca, New York.

Campuzano León, F. 2016. https://www.inaturalist.org/observations/12028134 (accessed 19 August 2024).

Castoe, T. A., J. M. Daza, E. N. Smith, M. M. Sasa, U. Kuch, J. A. Campbell, P. T. Chippindale, and C. L. Parkinson. 2009. Comparative phylogeography of pitvipers suggests a consensus of ancient Middle American highland biogeography. Journal of Biogeography 36:88–103.

Chambers, E. A., T. L. Marshall, and D. M. Hillis. 2023. The importance of contact zones for distinguishing interspecific from intraspecific geographic variation. Systematic Biology 72:357–371.

Chan, K. O., S. T. Hertwig, D. N. Neokleous, J. M. Flury, and R. M. Brown. 2022. Widely used, short 16S rRNA mitochondrial gene fragments yield poor and erratic results in phylogenetic estimation and species delimitation of amphibians. BMC Ecology and Evolution 22:1–9.

Chifman, J., and L. Kubatko. 2014. Quartet inference from SNP data under the coalescent model. Bioinformatics 30:3317–3324.

Crawford, A. J. 2003. Relative rates of nucleotide substitution in frogs. Journal of Molecular Evolution 57:636–641.

Daza, J.M., T. A. Castoe, and C. L. Parkinson. 2010. Using regional comparative phylogeographic data from snake lineages to infer historical processes in Middle America. Ecography 33:343–354.

de Queiroz, K. 1998. The general lineage concept of species, species criteria, and the process of speciation: a conceptual unification and terminological recommendations, p. 57–76. *In*: Endless Forms: Species and Speciation. D. J Howard, and S. H. Berlocher (eds.), Oxford Academic, New York, New York.

de Queiroz, K. 2005. A unified concept of species and its consequences for the future of taxonomy. Proceedings of the California Academy of Sciences 56:196–215.

de Queiroz, K. 2007. Species concepts and species delimitation. Systematic Biology 56:879–886.

de Queiroz, K. 2011. Branches in the lines of descent: Charles Darwin and the evolution of the species concept. Biological Journal of the Linnean Society 103:19–35.

Dell’Ampio, E., K. Meusemann, N. U. Szucsich, R. S. Peters, B. Meyer, J. Borner, M. Petersen, A. J. Aberer, A. Stamatakis, M. G. Walzl, B. Q. Minh, A. von Haeseler, I. Ebersberger, G. Pass, and B. Misof. 2014. Decisive data sets in phylogenomics: Lessons from studies on the phylogenetic relationships of primarily wingless insects. Molecular Biology and Evolution 31:239–249.

Denton, R. D., L. J. Kenyon, K. R. Greenwald, and H. L. Gibbs. 2014. Evolutionary basis of mitonuclear discordance between sister species of mole salamanders (*Ambystoma* sp.). Molecular Ecology 23:2811–2824.

Duellman, W. E. 1960. A distributional study of the amphibians of the Isthmus of Tehuantepec, Mexico. University of Kansas Publications, Museum of Natural History 13:19–72.

Duellman, W. E. 1970. The hylid frogs of Middle America. Monograph of the Museum of Natural History, the University of Kansas 1:1–753.

Dufresnes, C., D. Jablonski, J. Ambu, V.K. Prasad, K. Bala Gautam, R.G. Kamei, S. Mahony, S. Hofmann, R. Masroor, B. Alard, and A. Crottini. 2025. Speciation and historical invasions of the Asian black-spined toad (*Duttaphrynus melanostictus*). Nature Communications 16:298.

Durham, J. W., A. R. V. Arellano, and J. H. Peck. 1955. Evidence for no Cenozoic Isthmus of Tehuantepec seaways. Geological Society of America Bulletin 66:977–992.

Eaton, D.A., and I. Overcast. 2020. ipyrad: interactive assembly and analysis of RADseq datasets. Bioinformatics 36:2592–2594.

Evanno, G., S. Regnaut, and J. Goudet. 2005. Detecting the number of clusters of individuals using the software STRUCTURE: a simulation study. Molecular Ecology 14:2611–2620.

Ferguson, J. H., and C. H. Lowe, Jr. 1969. Evolutionary relationships in the *Bufo punctatus* group. American Midland Naturalist 81:435–466.

Ferrari, L., T. Orozco-Esquivel, V. Manea, and M. Manea. 2012. The dynamic history of the Trans-Mexican Volcanic Belt and the Mexico subduction zone. Tectonophysics 522:122–149.

Fox, J., and S. Weisberg. 2019. An R Companion to Applied Regression. Third edition. Sage, Thousand Oaks, California.

Frost, D. R. 2024. Amphibian Species of the World: an Online Reference. Version 6.2. https://amphibiansoftheworld.amnh.org/index.php (accessed 16 August 2024). American Museum of Natural History, New York, USA.

Frost, D. R., and A. G. Kluge. 1994. A consideration of epistemology in systematic biology, with special reference to species. Cladistics 10:259–294.

Frost, D. R., J. R. Mendelson III, and J. Pramuk. 2009. Further notes on the nomenclature of Middle American Toads (Bufonidae). Copeia 2009:418.

Frost, D. R., T. Grant, J. Faivovich, R. H. Bain, A. Haas, C. F. B. Haddad, R. O. de Sá, A. Channing, M. Wilkinson, S. C. Donnellan, C. J. Raxworthy, J. A. Campbell, B. L. Blotto, P. E. Moler … W. C. Wheeler. 2006. The amphibian tree of life. Bulletin of the American Museum of Natural History 297:1–370.

Garcia, R. A., M. Cabeza, C. Rahbek, and M. B. Araújo. 2014. Multiple dimensions of climate change and their implications for biodiversity. Science 344:1247579.

GBIF. 2024. GBIF Occurrence Download 10.15468/dl.5j329g (accessed 16 August 2024).

Girard, C. 1854. A list of North American bufonids, with diagnoses of new species. Proceedings of the Academy of Natural Sciences of Philadelphia 7:86–88.

Gloyd, H. K., and R. Conant. 1990. Snakes of the *Agkistrodon* complex—a monographic review. Contributions to Herpetology No. 6. Society for the Study of Amphibians and Reptiles, Oxford, Ohio.

Gorham, S. W. 1974. Checklist of World Amphibians Up to January 1, 1970. New Brunswick Museum, Saint John, Canada.

Günther, A. C. L. G. 1901. Reptilia and Batrachia, p. 237–252. *In*: Biologia Centrali Americana. O. Salvin, and F. D. Godman (eds.). London, R. H. Porter and Dulau & Co., London, England.

Halffter, G. 1987. Biogeography of the montane entomofauna of Mexico and Central America. Annual Review of Entomology 32:95–114.

Hardy D. K., F. X. González-Cózatl, E. Arellano, and D. S. Rogers. 2013. Molecular phylogenetics and phylogeographic structure of Sumichrast’s harvest mouse (*Reithrodontomys sumichrasti*: Cricetidae) based on mitochondrial and nuclear DNA sequences. Molecular Phylogenetics and Evolution 68:282–292.

Hillis, D. 2019. Species delimitation in herpetology. Journal of Herpetology 53:3–12.

Hoza, J., H. R. Davis, and A. D. Leaché. 2023. Genomic data do not support the species status of the San Luis Valley Short-Horned Lizard (*Phrynosoma diminutum*). Ichthyology & Herpetology 11:390–396.

Huidobro, L., J. J. Morrone, J. L. Villalobos, and F. Álvarez. 2006. Distributional patterns of freshwater taxa (fishes, crustaceans and plants) from the Mexican Transition Zone. Journal of Biogeography 33:731–741.

IUCN SSC Amphibian Specialist Group. 2020. Incilius marmoreus. IUCN Red List of Threatened Species 2020.

Jakobsson, M., and N. A. Rosenberg. 2007. CLUMPP: a cluster matching and permutation program for dealing with label switching and multimodality in analysis of population structure. Bioinformatics 23:1801–1806.

Johnson, B. B., T. A. White, C. A. Phillips, and K. R. Zamudio. 2015. Asymmetric introgression in a Spotted Salamander hybrid zone. Journal of Heredity 106:608–617

Lemos-Espinal, J. A., and J. R. Dixon. 2016. Anfibios y Reptiles de Hidalgo / Amphibians & Reptiles of Hidalgo. CONABIO, Ciudad de México, Distrito Federal, México.

Lemos-Espinal, J. A., G. R. Smith, and R. Valdes-Lares. 2019. Amphibians and Reptiles of Durango. ECO Herpetological Publishing & Distribution, Rodeo, New Mexico.

Liu, J-X., M-Y. Zhou, G-Q. Yang, Y-X. Zhang, P-F. Ma, C. Guo, M. S. Vorontsova, and D-Z. Li. 2020. ddRAD analyses reveal a credible phylogenetic relationship of the four main genera of *Bambusa*-*Dendrocalamus*-*Gigantochloa* complex (Poaceae: Bambusoideae). Molecular Phylogenetics and Evolution 146:106758.

Liu, S., M. Hou, Y-H. Lwin, Q. Wang, and D. Rao. 2021. A new species of *Gonyosoma* Wagler, 1828 (Serpentes, Colubridae), previously confused with *G. prasinum* (Blyth, 1854). Evolutionary Systematics 5:129–139.

Maldonado-Koerdell, M. 1964. Geohistory and paleogeography of Middle America, p. 3–32. In: Handbook of Middle American Indians, vol. 1. R.C. West (ed.). University of Texas Press, Austin, Texas.

Marshall T. L., E. A. Chambers, M. V. Matz, and D. M. Hillis. 2021. How mitonuclear discordance and geographic variation have confounded species boundaries in a widely studied snake. Molecular Phylogenetics and Evolution 162:107194.

Mastretta-Yanes, A., A. Moreno-Letelier, D. Piñero, T. H. Jorgensen, and B. C. Emerson. 2015. Biodiversity in the Mexican highlands and the interaction of geology, geography and climate within the Trans-Mexican Volcanic Belt. Journal of Biogeography 42:1586–1600.

Mateos, M. 2005. Comparative phylogeography of livebearing fishes in the genera *Poeciliopsis* and *Poecilia* (Poeciliidae: Cyprinodontiformes) in central Mexico. Journal of Biogeography 32:775–780.

Mayr, E. 1982. The Growth of Biological Thought: Diversity, Evolution, and Inheritance. Belknap Press of Harvard University Press, Cambridge, Massachusetts.

McCartney-Melstad, E., M. Gidiş, and H. B. Shaffer. 2018. Population genomic data reveal extreme geographic subdivision and novel conservation actions for the declining Foothill Yellow-legged Frog. Heredity 121:112–125.

Mendelson, J. R., III. 1994. A new species of toad (Anura: Bufonidae) from the lowlands of eastern Guatemala. Occasional Papers of the Museum of Natural History, University of Kansas 166:1–21.

Mendelson, J. R., III, D. G. Mulcahy, T. S. Williams, and J. W. Sites. 2011. A phylogeny and evolutionary natural history of Mesoamerican toads (Anura: Bufonidae: *Incilius*) based on morphology, life history, and molecular data. Zootaxa 3138:1–34.

Morrone, J. J., T. Escalante, and G. Rodríguez-Tapia. 2017. Mexican biogeographic provinces: map and shapefiles. Zootaxa 4277:277–279.

Morrone, J.J. 2020. Biogeographic regionalization of the Mexican Transition Zone, p. 103. In: The Mexican Transition Zone. Springer, Cham, Switzerland.

Mulcahy, D. G., B. H. Morrill, and J.R. Mendelson, III. 2006, Historical biogeography of lowland species of toads (*Bufo*) across the Trans-Mexican Neovolcanic Belt and the Isthmus of Tehuantepec. Journal of Biogeography 33:1889–1904.

Myers, E. A., A. D. McKelvy, and F. T. Burbrink. 2020. Biogeographic barriers, Pleistocene refugia, and climatic gradients in the southeastern Nearctic drive diversification in cornsnakes (*Pantherophis guttatus* complex). Molecular Ecology 29:797–811.

Oliveira, B. F., V. A. São-Pedro, G. Santos-Barrera, C. Penone, and G. C. Costa. 2017. AmphiBIO, a global database for amphibian ecological traits. Scientific Data 4:1–7.

Pante, E., J. Abdelkrim, A. Viricel, D. Gey, S. C. France, M. C. Boisselier, and S. Samadi. 2015. Use of RAD sequencing for delimiting species. Heredity 114:450–459.

Pereyra, M. O., B. J. Blotto, D. Baldo, J. C. Chaparro, S. R. Ron, A. J. Elias-Costa, P. P. Iglesias, P. J. Venegas, M. T. C. Thome, J. J. Ospina-Sarria, and N. M. Maciel. 2021. Evolution in the genus *Rhinella*: a total evidence phylogenetic analysis of neotropical true toads (Anura: Bufonidae). Bulletin of the American Museum of Natural History 447:1–156.

Peterson, B. K., J. N. Weber, E. H. Kay, H. S. Fisher, and H. E. Hoekstra. 2012. Double digest RADseq: an inexpensive method for de novo SNP discovery and genotyping in model and non-model species. PLOS ONE 7:e37135.

Porras, L. W., L. D. Wilson, G. W. Schuett, and R. S. Reiserer. 2013. A taxonomic reevaluation and conservation assessment of the Common Cantil, *Agkistrodon bilineatus* (Squamata: Viperidae): a race against time. Amphibian and Reptile Conservation 7:48–73.

Pritchard, J. K., M. Stephens, and P. Donnelly. 2000. Inference of population structure using multilocus genotype data. Genetics 155:945–959.

Pyron R. A., K. A. O’Connell, E. M. Lemmon, A. R. Lemmon, and D. A. Beamer. 2020. Phylogenomic data reveal reticulation and incongruence among mitochondrial candidate species in Dusky Salamanders (*Desmognathus*). Molecular Phylogenetics and Evolution 146:106751.

Rambaut, A., A. J. Drummond, D. Xie, G. Baele, and M. A. Suchard. 2018. Posterior summarization in Bayesian phylogenetics Using Tracer 1.7. Systematic Biology 67:901–904.

Rannala, B., and Z. Yang. 2003. Bayes estimation of species divergence times and ancestral population sizes using DNA sequences from multiple loci. Genetics 164:1645–1656.

Ríos-Muñoz, C.A., and A.G Navarro□Sigüenza. 2012. Patterns of species richness and biogeographic regionalization of the avifaunas of the seasonally dry tropical forest in Mesoamerica. Studies on Neotropical Fauna and Environment 47:171–182.

Rivera, D., I. Prates, T.J. Firneno, M. Trefaut Rodrigues, J.P. Caldwell, and M.K. Fujita. 2022. Phylogenomics, introgression, and demographic history of South American true toads (*Rhinella*). Molecular Ecology 31:978–992.

Ruane, S., R. W. Bryson, R. A. Pyron, and F. T. Burbrink. 2014. Coalescent species delimitation in Milksnakes (Genus *Lampropeltis*) and impacts on phylogenetic comparative analyses. Systematic Biology 63:231–250.

Rzedowski, J. 1994. Vegetación de México. Limusa Noriega Editores, Ciudad de México, Distrito Federal, México.

Sabaj, M. H. 2023. Codes for Natural History Collections in Ichthyology and Herpetology (online supplement). Version 9.5 (10 Nov 2023). Electronically accessible at https://asih.org, American Society of Ichthyologists and Herpetologists, Washington, DC.

Sanders, O., and H. M. Smith. 1951. Geographic variation in toads of the *debilis* group of *Bufo*. Field and Laboratory 19:141–160.

Schliep, K. P. 2011. phangorn: phylogenetic analysis in R. Bioinformatics 27:592–593.

Schliep, K., A. J. Potts, D. A. Morrison, and G. W. Grimm. 2017. Intertwining phylogenetic trees and networks. Methods in Ecology and Evolution 8:1212–1220.

Schramm, F. D., A. Valdez-Mondragón, and L. Prendini. 2021. Volcanism and palaeoclimate change drive diversification of the world’s largest whip spider (*Amblypygi*). Molecular Ecology 30:2872–2890.

Scott, P. A., L. J. Allison, K. J. Field, R. C. Averill-Murray, and H. B. Shaffer. 2020. Individual heterozygosity predicts translocation success in threatened desert tortoises. Science 370:1086–1089.

Smith, H. M., and D. Chiszar. 2001. A new subspecies of Cantil (*Agkistrodon bilineatus*) from central Veracruz, Mexico (Reptilia: Serpentes). Bulletin of the Maryland Herpetological Society 37:130–136.

Stamatakis, A. 2014. RAxML version 8: a tool for phylogenetic analysis and post-analysis of large phylogenies. Bioinformatics 30:1312–1313.

Suárez□Atilano, M., F. Burbrink, and E. Vázquez□Domínguez. 2014. Phylogeographical structure within *Boa constrictor imperator* across the lowlands and mountains of Central America and Mexico. Journal of Biogeography 41:2371–2384.

Suazo-Ortuño, I., A. Ramírez-Bautista, and J. Alvarado-Díaz. 2023. Amphibians and reptiles of Mexico: diversity and conservation, p. 105–125. In: Mexican Fauna in the Anthropocene. R. W. Jones, C. P. Ornelas-García, R. Pineda-López, and F. Álvarez. (eds.). Springer, Cham, Switzerland.

Sullivan, J., E. Arellano, and D. S. Rogers. 2000. Comparative phylogeography of Mesoamerican highland rodents: concerted versus independent response to past climatic fluctuations. The American Naturalist 155:755–768.

Sullivan, J., J. A. Markert, and C. W. Kilpatrick. 1997. Phylogeography and molecular systematics of the *Peromyscus aztecus* species group (Rodentia: Muridae) inferred using parsimony and likelihood. Systematic Biology 46:426–40.

Sun, Y-B., Z-J. Xiong, X-Y. Xiang, S-P. Liu, W-W. Zhou, X-L. Tu, L. Zhong, L. Wang, D-D. Wu, B-L. Zhang, C-L. Zhu, M-M. Yang, H-M. Chen, F. Li … Y-P. Zhang. 2015. Whole-genome sequence of the Tibetan frog *Nanorana parkeri* and the comparative evolution of tetrapod genomes. Proceedings of the National Academy of Sciences 112:1257–1262.

Taylor, E. H. 1940 “1939”. Herpetological miscellany No. I. University of Kansas Science Bulletin 26:489–571.

Toews, D. P. L., and A. Brelsford. 2012. The biogeography of mitochondrial and nuclear discordance in animals. Molecular Ecology 21:3907–3930.

van der Heiden, A. 2022. *Incilius marmoreus* (Marbled Toad). Sexual dichromatism and reproductive behavior. Herpetological Review 53:468–470.

Wall, M. E., A. Rechtsteiner, and L. M. Rocha. 2003. Singular value decomposition and principal component analysis, p. 91–109. In: A Practical Approach to Microarray Data Analysis. D. P. Berrar, W. Dubitzky, M. Granzow. (eds.). Springer, Boston, Massachusetts.

Wang, K., J. A. Lenstra, L. Liu, Q. Hu, T. Ma, Q. Qiu, and J. Liu. 2018. Incomplete lineage sorting rather than hybridization explains the inconsistent phylogeny of the wisent. Communications Biology 1:169.

Weir, J. J., E. Bermingham, M. J. Miller, J. Klicka, and M. A. González. 2008. Phylogeogoraphy of a morphologically diverse Neotropical montane species, the Common Bush-Tanager (*Chlorospingus ophthalmicus*). Molecular Phylogenetics and Evolution 47:650–64.

Whitney, K. D., J. R. Ahern, L. G. Campbell, L. P. Albert, and M. S. King. 2010. Patterns of hybridization in plants. Perspectives in Plant Ecology, Evolution and Systematics 12:175–182.

Wickham, H. 2016. ggplot2: Elegant Graphics for Data Analysis. Springer-Verlag, New York, New York.

Yang, Z. 2015. The BPP program for species tree estimation and species delimitation. Current Zoology 61:854–865.

Zamudio-Beltrán, L. E., Y. Licona-Vera, B. E. Hernández-Baños, J. Klicka, and J. F. Ornelas. 2020. Phylogeography of the widespread white-eared hummingbird (*Hylocharis leucotis*): pre-glacial expansion and genetic differentiation of populations separated by the Isthmus of Tehuantepec. Biological Journal of the Linnean Society 130:247–267.

Zarza, E., V. H. Reynoso, C. M. Faria, and B. C. Emerson. 2019. Introgressive hybridization in a Spiny-Tailed Iguana, *Ctenosaura pectinata*, and its implications for taxonomy and conservation. PeerJ 7:e6744.

