## Supplementary material for "Resolving the Taxonomic Status of the Marbled Toad (Bufonidae: *Incilius marmoreus*): 2RAD-based Phylogeography Including an Isolated Population in Veracruz, Mexico": Figure 1 Alt-Text

**A two-panel figure containing a map of Mexico showing current range of *Incilius marmoreus* with locality origin of tissues used (A, left) joined by a Principal Components Analysis (PCA) scatterplot of 2RAD data (B, right).**

**Panel A (left):** A shaded elevational map of Mexico, wherein darker shades of gray confer higher elevation (white = 0 meters, dark gray = above 2400 meters). Small white dots located along the Pacific Coast and near the coastal city of Veracruz represent iNaturalist reported sightings of *Incilius marmoreus* as confirmed by the authors. Localities of tissues used for 2RAD sequencing are plotted as larger symbols. Different shapes and colors denote different states: Veracruz (one black circle representing one distinct locality), Oaxaca (two yellow downward triangles representing two distinct localities), Guerrero (six light blue upward triangles), Colima (one green diamond), Jalisco (one dark blue square), Nayarit (one dotted orange square), and Sinaloa (one asterisked pink square). These symbols remain consistent throughout all figures used in this paper.

**Panel B (right):** PCA scatterplot showing variation of 2RAD data between *Incilius marmoreus* populations. The x-axis is PC1 (31.1% of variation) and the y-axis is PC2 (12.4% of variation). Different shapes and colors denote different states: Veracruz (two black circles representing two samples), Oaxaca (four yellow downward triangles representing two samples), Guerrero (six light blue upward triangles), Colima (one green diamond), Jalisco (one dark blue square), Nayarit (one dotted orange square), and Sinaloa (one asterisked pink square). Samples from Veracruz cluster at the top left, samples from Oaxaca occupy a swath below those from Veracruz samples, and samples from Guerrero samples cluster below those from Oaxaca. The sample from Colima is to the right of the aforementioned localities, near the bottom and about a quarter-way to the right of the y-axis. The sample from Sinaloa is located considerably to the right of all the aforementioned localities, and below and to the right are the single samples from Nayarit and Jalisco; the latter two samples overlap considerably.
