## Supplementary material for "Resolving the Taxonomic Status of the Marbled Toad (Bufonidae: *Incilius marmoreus*): 2RAD-based Phylogeography Including an Isolated Population in Veracruz, Mexico": Figure 2 Alt-Text

**A two-panel figure displaying Principal Component Analysis (PCA) results based on morphological measurements.**

**Panel A (left):** PCA of male toads with PC1 (57.0%) on the x-axis and PC2 (14.9%) on the y-axis. Symbols: Veracruz (black circles), Chiapas (gray circles), Oaxaca (yellow downward triangles), Guerrero (light blue upward triangles), Colima (green diamonds), Jalisco (dark blue squares), Nayarit (orange dotted squares), Sinaloa (pink asterisked squares), and Michoacán (yellow dotted circles). Ellipses indicate 95% confidence intervals: a solid black ellipse encloses the samples from Veracruz and a dashed ellipse encloses all other samples. While localities display clustering patterns, there are significant overlaps. Notably, the ellipses representing samples from Veracruz and ‘all others” overlap considerably, though some samples from Veracruz are located particularly to the center-lower right.

**Panel B (right):** PCA of female toads with PC1 (59.2%) on the x-axis and PC2 (12.2%) on the y-axis, using the same color and shape coding. Symbols: Veracruz (black circles), Chiapas (gray circles), Oaxaca (yellow downward triangles), Guerrero (light blue upward triangles), Colima (green diamonds), Jalisco (dark blue squares), Nayarit (orange dotted squares), Sinaloa (pink asterisked squares), and Michoacán (yellow dotted circles). Ellipses indicate 95% confidence intervals: a solid black ellipse encloses the samples from Veracruz and a dashed ellipse encloses all other samples. Likewise, while localities display clustering patterns, there are significant overlaps. Notably, the ellipses representing samples from Veracruz and “all others” overlap considerably, though some samples from Veracruz are located particularly to the center-upper left.
