## Supplementary material for "Resolving the Taxonomic Status of the Marbled Toad (Bufonidae: *Incilius marmoreus*): 2RAD-based Phylogeography Including an Isolated Population in Veracruz, Mexico": Figure 3 Alt-Text

A three-panel figure of phylogenetic plots based on 2RAD sequenced data. Representation of symbols in all panels: Veracruz (black circles), Oaxaca (yellow downward triangles), Guerrero (light blue upward triangles), Colima (green diamonds), Jalisco (dark blue squares), Nayarit (orange dotted squares), and Sinaloa (pink asterisked squares).

**Panel A (left):** DensiTree-like plot of RAxML rapid bootstrap replicates from a maximum-likelihood analysis of loci from 2RAD data, rooted with *Rhinella horribilis*. Thicker black lines indicate stronger concordance among replicates. The topology shows *I. marmoreus* and *I. perplexus* as sister groups, with that pair being sister to *I. canaliferus*. *Incilius mazatlanensis* and *I. valliceps* are sister to each other and are together sister to the aforementioned trio of taxa. Within *I. marmoreus*, the topology shows samples from Veracruz as nested within those from Oaxaca and otherwise most closely related to samples from Guerrero. These samples collectively are then most closely related to the single sample from Colima and then to a clade consisting of the samples from Jalisco, Nayarit, and Sinaloa.

**Panel B (center):** Similar DensiTree-like plot showing bootstrap replicates from a tetrad species-tree analysis based on SNP data from the same dataset. Topology is consistent with panel A.

**Panel C (right):** Unrooted Neighbor-Net phylogenetic network inferred from one random SNP per locus, showing relationships between Incilius marmoreus and related species. Incilius marmoreus populations cluster together, with populations from Guerrero, Oaxaca and Veracruz most similar to each other—similar to Panels A and B. Samples from all other species are more distant.
