## Supplementary material for "Resolving the Taxonomic Status of the Marbled Toad (Bufonidae: *Incilius marmoreus*): 2RAD-based Phylogeography Including an Isolated Population in Veracruz, Mexico": Figure 4 Alt-Text

A two-panel figure of shaded elevational maps of Mexico displaying the genetic clustering results for *Incilius marmoreus*, based on SNP data from 2RAD and analyzed with the program Structure. Darker shades of gray confer higher elevation (white = 0 meters, dark gray = above 2400 meters).

Panel A: (k = 2), thirteen pie charts colored in varying proportions of blue (darker gray) and yellow (lighter gray) scattered along the Pacific Coast and one in Veracruz, illustrating the genetic cluster assignment of different individuals from Structure analyses. Each pie chart represents the average assignment probabilities from each site. Three pie charts along the northern Pacific Coast are almost entirely blue (darker gray). Nine pie charts including the southern Pacific Coast and Veracruz are almost entirely yellow (lighter gray). One pie chart near the town of Manzanillo, in northwestern Colima, is about twenty percent blue (lighter gray) and eighty percent yellow (darker gray).

Panel B: (k = 4) shows the same map with a finer genetic division into four clusters, represented by thirteen pie charts colored in varying proportions of pink (asterisked), blue (darker gray), yellow (lighter gray), and black. Pie charts are described moving southward along the Pacific Coast, then the population in Veracruz. One pie chart just south of Culiacán, Sinaloa, is almost entirely pink (asterisked). Two pie charts near Puerto Vallarta, Jalisco, are almost entirely blue (darker gray). The pie chart near the town of Manzanillo, northwestern Colima, is about three-quarters yellow (lighter gray), with the remainder split roughly equally between blue (darker gray) and black. In the state of Guerrero, five pie charts are almost entirely yellow (lighter gray), and the southernmost pie chart is about ten percent black. This grade continues southward into the state of Oaxaca, where two pie charts contain about seventy to eighty percent yellow (lighter gray), with the remainder black. In Veracruz, one pie chart is about seventy percent black, with the remainder yellow (lighter gray).
