## Supplementary material for "Resolving the Taxonomic Status of the Marbled Toad (Bufonidae: *Incilius marmoreus*): 2RAD-based Phylogeography Including an Isolated Population in Veracruz, Mexico": Figure 5 Alt-Text

A two-panel figure showing results from BPP coalescent analyses with two demographic scenarios.

Panel A (left): represents the coalescence between samples from Veracruz and all other populations. On the left, the bar for “all other populations” is blue and thick, proportionately corresponding to the effective population size of N_e_ = 0.81M (million). On the right, the bar for Veracruz is thin and black, with an estimated N_e_ of 0.04M. The Veracruz and “all other populations” bars have a height corresponding to a coalescence date of 0.33 million years ago (Mya), with a 95% confidence interval between 0.19 and 0.56 Mya. In between and above the Veracruz and “all other populations” bar is the ancestral population, depicted by a thick gray bar. Its effective population size (N_e_) is 0.85M.

Panel B (right): represents a scenario where coalescence is estimated separately for samples from Veracruz and the samples from the southern Pacific Coast, and then for both of these with the samples from the northern Pacific Coast. The bar for “southern” is thick and yellow, with N_e_ = 0.33M. It shares the same height as the thin and black bar for “Veracruz”, which has N_e_ = 0.04M, representing a coalescent timing of 0.31 Mya (interval: 0.17 to 0.53 Mya). The ancestral population giving rise to the southern and Veracruz populations is represented as a moderately thick light gray bar in between and above the southern and Veracruz bars with N_e_ = 0.19M. To the left of these bars is the “northern” population, a moderately thick bar with N_e_ = 0.28M. Its height represents the coalescent date (0.86 Mya, with an interval: 0.51 to 1.41 Mya) between this northern population and the ancestral population giving rise to the southern and Veracruz populations. Finally, located in the center and above all other bars, a very thick and dark gray bar represents the ancestral population giving rise to the northern population and the ancestral population of the southern and populations in Veracruz. It has a N_e_ = 0.95M.
