## Supplementary material for "Resolving the Taxonomic Status of the Marbled Toad (Bufonidae: *Incilius marmoreus*): 2RAD-based Phylogeography Including an Isolated Population in Veracruz, Mexico": Table 1 Summary

Table 1: This table presents linear model output statistics determined from a series of linear models for morphological measurements of male Marbled Toads (*Incilius marmoreus*). The first column labels the measurement type (TIB = tibial length; FT =foot length; HL = head length; HW = head width; EYE = eye diameter; EN = eye-nostril length;

TYMP = tympanum diameter; PARL = parotoid length; PARW = parotoid width) and whether the reported significance values concern tests on slopes or intercepts. The second column presents type three Sum of Squares values associated with the measurement types denoted in the first column. The third column presents type two Sum of Squares values. The fourth column presents F-values. The fifth column presents p-values. Statistically significant p-values

(α = 0.05) are marked with an asterisk and values statistically significant after a Bonferroni correction (adjusted α =0.0056) are marked with two asterisks. Head width slopes are significant (p = 0.034), as are eye diameter slopes (p = 0.045). Eye-nostril length slopes are significant after Bonferroni correction (p = 0.000), as are tympanum diameter intercepts (p = 0.004).
