## Supplementary material for "Resolving the Taxonomic Status of the Marbled Toad (Bufonidae: *Incilius marmoreus*): 2RAD-based Phylogeography Including an Isolated Population in Veracruz, Mexico": Table 3 Summary

Table 3: This table presents results from principal components analysis for male Marbled Toads (Incilius marmoreus). The first column reads from top to bottom: “Proportion of Variance, Cumulative Variance, Eigenvalue, SVL (snout-vent length), TIB (tibial length), FT (foot length), HL (head length), HW (head width), EYE (eye diameter), EN (eye-nostril length), TYMP (tympanum diameter), PARL (parotoid length), PARW (parotoid width). The second column presents Proportion of Variance, Cumulative Variance, Eigenvalue, and loadings for the first principal components axis. The third column presents similar statistics and loadings for the second axis. The fourth column presents similar statistics and loadings for the third axis.
